# The neural basis of tadpole transport in poison frogs

**DOI:** 10.1101/630681

**Authors:** Eva K. Fischer, Alexandre B. Roland, Nora A. Moskowitz, Elicio E. Tapia, Kyle Summers, Luis A. Coloma, Lauren A. O’Connell

**Affiliations:** Department of Biology, Stanford University, Stanford, California, United States of America; Center for Systems Biology, Harvard University, Cambridge, Massachusetts, United States of America; Centro Jambatu de Investigación y Conservación de Anfibios, Fundación Otonga, Quito, Ecuador; Department of Biology, East Carolina University, Greenville, North Carolina, United States of America

**Keywords:** parental care, poison frog, phosphoTRAP, preoptic area, hippocampus, galanin

## Abstract

Parental care has evolved repeatedly and independently across animals. While the ecological and evolutionary significance of parental behaviour is well recognized, underlying mechanisms remain poorly understood. We took advantage of behavioural diversity across closely related species of South American poison frogs (Family Dendrobatidae) to identify neural correlates of parental behaviour shared across sexes and species. We characterized differences in neural induction, gene expression in active neurons, and activity of specific neuronal types in three species with distinct parental care patterns: male uniparental, female uniparental, and biparental. We identified the medial pallium and preoptic area as core brain regions associated with parental care, independent of sex and species. Identification of neurons active during parental care confirms a role for neuropeptides associated with care in other vertebrates as well as identifying novel candidates. Our work is the first to explore neural and molecular mechanisms of parental care in amphibians and highlights the potential for mechanistic studies in closely related but behaviourally variable species to build a more complete understanding of how shared principles and species-specific diversity govern parental care and other social behaviour.

## Background

Parental care is an important adaptation that allows exploitation of novel habitats, influences fitness and survival of parents and offspring, and serves as an evolutionary precursor to other affiliative behaviour [1,2]. Specialized parental care strategies have evolved repeatedly and independently across animals, yet the mechanisms underlying parental behaviour and its evolution remain poorly understood. The neural mechanisms promoting parental care in females are best understood in mammals [3]. However, female uniparental care evolved at the base of the mammalian lineage and therefore provides limited clues to the evolutionary origins of parenting. Moreover, studies of male parental care come mostly from biparental systems [4,5] in which parental behaviour cannot easily be dissociated from pair bonding. What is needed to further understand the mechanisms underlying parental behaviour and its evolution are studies across closely-related species that vary in care strategies.

Parental care can be conceptualized as a complex set of inter-related behaviours controlled by brain regions involved in the integration of sensory, social, motivational, and cognitive aspects of care [6]. Across vertebrates, many of these functions are performed by the social decision-making network (SDMN; [7]), a highly interconnected group of evolutionarily ancient and functionally conserved brain regions. Although studies on the neural mechanisms of parental behaviour are sparse outside mammals, and particularly lacking in amphibians and reptiles, the SDMN provides an ideal starting point for this work as network nodes and connectivity are well understood and highly conserved, and behaviourally important ligand/receptor complexes have been extensively studied [7,8].

Dendrobatid poison frogs show remarkable diversity in parental care across closely related species, including male uniparental, female uniparental, and biparental care. Parental care in poison frogs involves egg attendance during embryo development, generally followed by transportation of tadpoles “piggyback” to pools of water upon hatching [9–11]. In some species, mothers nourish growing tadpoles with unfertilized, trophic eggs until metamorphosis [10–12].

Importantly, both male and female care occur with and without pair bonding in this clade [13], allowing the dissociation of pair bonding from parental care. The diversity of behavioural care strategies among poison frog species affords a unique opportunity to identify physiological, neural, and molecular contributions to parental care across sexes of closely related species as well as across the convergent evolution of parental behaviour among major vertebrate lineages.

In the current study, we take advantage of three focal species with distinct care patterns: *Dendrobates tinctorius* (male uniparental), *Ranitomeya imitator* (biparental and monogamous), and *Oophaga sylvatica* (female uniparental). By comparing neural activity in parental frogs and their non-caregiving reproductive partners, we identify core brain regions active during tadpole transport independent of sex and species. To identify neuronal types mediating tadpole transport, we characterize gene expression and activity patterns specifically in behaviourally relevant neurons within core brain regions. Our experiments are the first to explore neural and molecular mechanisms of parental care in amphibians and demonstrate the utility of mechanistic studies in closely related, behaviourally distinct species to identify core neural correlates of parental behaviour.

## Methods

### Laboratory sample collection

*Dendrobates tinctorius* and *Ranitomeya imitator* frogs were housed in breeding pairs in the laboratory, allowing us to identify both parental individuals and their non-caregiving reproductive partners. To control for effects of experience, all pairs successfully reared at least one clutch from egg-laying through tadpole transport prior to the experiment. For the non-parental group, we collected frog pairs between parental bouts when they were not caring for eggs or tadpoles, collecting individuals of both the caregiving sex (non-transport; n=10 *D. tinctorius*, n=7 *R. imitator*) and their opposite sex reproductive partners (non-transport partner; n=9 *D. tinctorius*, n=8 *R. imitator*). For the tadpole transport group, when we found transporting frogs, we collected both the transporting individual (tadpole transporter; n=13 *D. tinctorius*, n=7 *R. imitator*) and its opposite sex, non-caregiving partner (transport partner; n=11 *D. tinctorius*, n=6 *R. imitator*). All brain tissue was collected in an identical manner: frogs were captured, anesthetized with benzocaine gel, weighed and measured, and euthanized by rapid decapitation. This entire process took less than 5 minutes. All procedures were approved by the Harvard University Animal Care and Use Committee (protocol no. 12-10-1).

### Field sample collection

*Oophaga sylvatica* (Puerto Quito-Santo Domingo population) were collected in field enclosures in Ecuador in April and May of 2016. We collected non-parental control females (N=8) from enclosures containing only mature females to ensure that frogs were not currently caring for eggs or tadpoles. We collected tadpole transporting females (N=5) from enclosures containing multiple males and females and therefore could not identify their non-caregiving male reproductive partners. Frogs were captured, anesthetized with benzocaine gel, weighed and measured, and euthanized by rapid decapitation. Procedures were approved by the Harvard University Animal Care and Use Committee (protocol no. 15-03-239). All samples were collected and imported in accordance with Ecuadorian and US Law (collection permits: 005-15-IC-FAU-DNB/MA and 007-2016-IC-FAU-DNB/MA; CITES export permit 16EC000007/VS issued by the Ministerio de Ambiente de Ecuador).

### Immunohistochemistry

Whole brains were placed into 4% paraformaldehyde at 4°C overnight and then transferred to a 30% sucrose solution for cryoprotection. Once dehydrated, brains were embedded in Tissue-Tek^®^ O.C.T. Compound (Electron Microscopy Sciences, Hatfield, PA, USA), rapidly frozen, and stored at −80°C until cryosectioning. We sectioned brains into four coronal series at 14μm, allowed slides to dry completely, and stored slides at −80°C.

To assess the level of neural activity across brain regions, we used an antibody for phosphorylated ribosomes (pS6; phospho-S6 Ser235/236; Cell Signaling, Danvers, MA, USA) and followed standard immunohistochemical procedures for 3’,3’-diaminobenzadine (DAB) antibody staining (as in [14]). To ask whether neural activity was higher specifically in galanin neurons, we combined the pS6 antibody with a custom-made galanin antibody (peptide sequence: CGWTLNSAGYLLGPHAVDNHRSFNDKHGLA; Pocono Rabbit Farm & Laboratory, Inc, Canadensis, PA, USA) and followed standard immunohistochemical procedures for fluorescent double labeling (as in [4]). Additional methodological details are in Supplemental Materials.

### Microscopy and cell counts

Stained brain sections were photographed on a Leica DMRE connected to a QImaging Retiga 2000R camera at 20X magnification. We quantified labeled cells from photographs using FIJI image analysis software [15]. Brain regions were identified using a custom dendrobatid frog brain atlas (Supplemental Materials). We measured the area of candidate brain regions and counted all labeled cells in a single hemisphere for each brain region across multiple sections. We quantified cell number in the nucleus accumbens, the basolateral nucleus of the stria terminalis, the habenula, the lateral septum, the magnocellular preoptic area, the medial pallium (homolog of the mammalian hippocampus), the anterior preoptic area, the suprachiasmatic nucleus, the striatum, the posterior tuberculum (homolog of the mammalian midbrain dopamine cells representing the ventral tegmental area and substantia nigra), the ventral hypothalamus, and the ventral pallium.

Fluorescently stained brain sections were photographed at 20X magnification on a Leica DM4B compound microscope attached to a fluorescent light source. Each section was visualized at three wavelengths (594nm, 488nm, 358nm) and images were pseudo-coloured to reflect these spectra. We used DAPI nuclear staining to identify brain regions as above and quantified the number of galanin positive cells, pS6 positive cells, and co-labeled cells from photographs of the preoptic area using FIJI image analysis software [15]. We combined counts for all preoptic area sub-regions due to the low overall number of galanin-positive neurons and because this more closely reflected the neuroanatomical resolution of tissue punches used in PhophoTRAP (see below).

### Statistical analyses of cell counts

We analyzed the relationship between parental behaviour and pS6 neural activity to identify brain regions whose activity differed during tadpole transport independent of sex and species (i.e. core parental care brain regions). We used generalized linear mixed models with a negative binomial distribution appropriate for count data with unequal variances to test for differences in pS6 positive cell number. For laboratory animals, behavioural group (tadpole transport vs non-parental), sex, brain region, and their interactions were included as main effects predicting the number of pS6-positive cells. For field sampled *O. sylvatica*, sex was omitted from the model as we could not identify non-caregiving reproductive partners and collected only females. Rather than averaging cell counts across brain regions, we included frog identity as a repeated measure to control for both random and systematic variation across the large number of tissue sections quantified for each individual. Brain region area was included as an offset variable to control for body size differences between frogs, known size differences between brain regions, and rostral to caudal size/shape variation within brain regions. We explored main effects of group, sex, and regional differences in further detail using *post hoc* comparisons Tukey adjusted for multiple hypothesis testing.

We tested for differences in the number and activity of galanin neurons using generalized linear mixed models. To compare the number of galanin neurons, we included behavioural group (tadpole transport vs non-parental), sex, and their interactions as main effects predicting the median number of galanin positive cells using a negative binomial distribution appropriate for count data with unequal variances. To analyze activity differences in preoptic area galanin neurons, we included behavioural group, sex, and their interactions as main effects predicting the proportion of pS6 positive galanin (i.e. co-labeled) cells using a binomial distribution. All analyses were performed separately for each species using SAS Statistical Software (SAS 9.4; SAS Institute for Advanced Analytics) and representative SAS code is provided in the Supplemental Materials.

### PhosphoTRAP library construction & sequencing

We collected *D. tinctorius* males found transporting tadpoles and males that currently had tadpoles present in the leaf litter but had not yet transported them. Males were euthanized as described above (N=9 per group). Brains were removed, embedded in Tissue-Tek^®^ O.C.T. Compound, frozen on dry ice, and stored at −80°C for no more than 1 month. Once all animals had been collected, brains were sectioned at 100 μm on a cryostat and thaw mounted onto SuperFrost Plus slides. A 0.96 mm tissue micro punch tool was used to isolate the medial pallium and rostral hypothalamus (anterior, medial, and magnocellular preoptic area and suprachiasmatic nucleus). To provide enough starting material for PhosphoTRAP, brain regions from three individuals were combined into a single sample, for a total of three biological replicates per group. PhosphoTRAP libraries for total (TOT) and immunoprecipitated (IP) RNA from each sample were constructed following [16] (details in Supplemental Materials). Libraries were then pooled in equimolar amounts and sequenced on an Illumina HiSeq 2500.

### PhosphoTRAP analysis

To analyze PhosphoTRAP data we first quantified gene expression by mapping sequenced reads back to a brain tissue specific *D. tinctorius* transcriptome (Fischer & O’Connell, *unpublished)* and estimated their abundance using Kallisto [17]. As gene expression is known to differ across brain regions [18], we performed all subsequent analysis steps separately for the Mp and POA. Analysis methods are described in detail in the Supplemental Materials. Briefly, we normalized read counts using DESeq2 [19] and quantified transcript enrichment/depletion in active neurons as a log-fold difference between transcript counts from immunoprecipitated (IP) and total (TOT) mRNA for each sample. We then calculated differential fold enrichment between parental and non-parental individuals by dividing mean log-fold expression values from the two behavioural groups. We refer to this final metric as the log-fold difference ratio between tadpole transport and non-transport behavioural groups.

Our primary objective was to utilize PhosphoTRAP data to identify Mp and POA cell types whose activity differed between tadpole transport and non-parental individuals. To this end, we restricted further analysis to a subset of 158 transcripts representing cell types with known roles in parental care (Table S1). We identified transcripts as significantly enriched/depleted based on a combination of log-fold enrichment thresholds (>4) and permutation testing (Supplemental Materials). Permutation testing and visualization were done using R Statistical Software (version 3.5.0; the R Foundation for Statistical Computing).

## Results

### Neural induction during tadpole transport

We compared neural activity patterns in tadpole transporters and their non-transporting reproductive partners across three closely related poison frog species with distinct parental care strategies (Fig. 1A). Differences in neural activity depended on behavioural group, sex, and brain region (Fig. 1B; Table 1) and associations between behavioural group and neural induction were brain region specific (Table 1; group*region: *D. tinctorius*: F_1,2515_=5.00, p<0.0001; *R. imitator*. F_12,557_=6.85, p<0.0001; *O. sylvatica*: F_12,557_=5.53, p<0.0001). We found overall differences between the transporting and non-transporting sex in male uniparental *D. tinctorius* (sex*group*region: F_1,2515_=3.89, p<0.0001) but not biparental *R. imitator* (Table 1; Fig. 1B). Indeed, *post hoc* analyses of region-specific differences revealed greater similarity between sexes in biparental, monogamous *R. imitator* than male uniparental *D. tinctorius* (Fig. 1B,C; Table S2).

**Figure 1.**
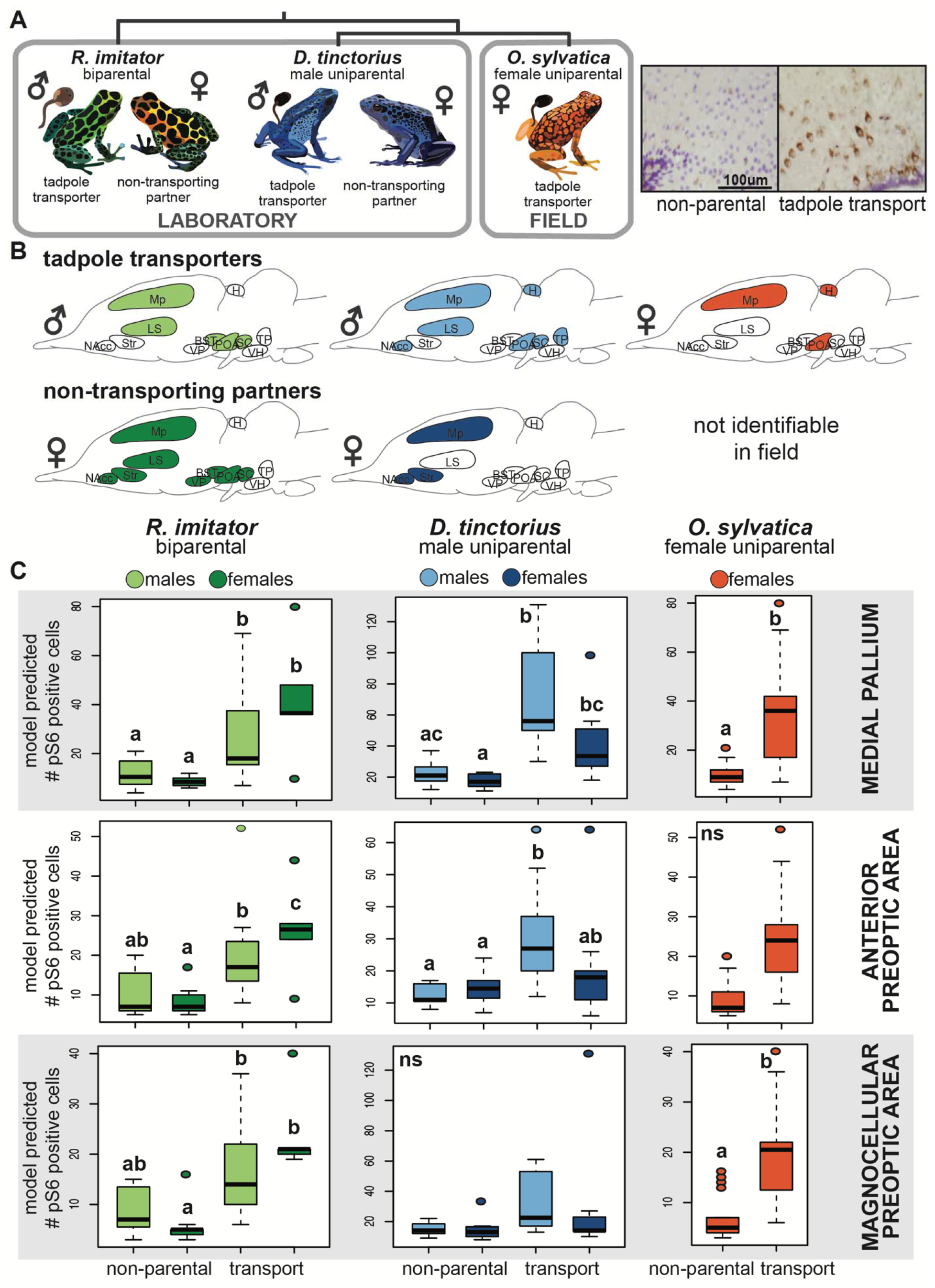
Patterns of neural induction associated with parental care. **(A)** Overview of experimental design. Our comparative approach allowed us to identify brain regions important in parental care independent of sex and species. **(B)** Overview of brain regions showing differences in neural activity between parental and non-parental individuals (shaded) for tadpole transporting sex and their non-transporting partners. Small symbols indicate the sex of transporting and non-transporting partner individuals. Comparing across species, we identified Mp and POA as active during tadpole transport regardless of sex and species (i.e. as core parental care brain regions). **(C)** Detailed results for core brain regions. Letters above the box plots indicate statistical differences: comparing groups pairwise, shared letters indicate no significant differences, non-shared letters indicate significant differences (p<0.05). Representative micrographs of pS6 staining (brown) with cresyl violet nuclear stain (purple) from the mPOA are shown at top right. Abbreviations: BST = basolateral nucleus of the stria terminalis, H = habenula, Ls = lateral septum, Mp = medial pallium (homolog of the mammalian hippocampus), NAcc = nucleus accumbens, aPOA = anterior preoptic area, mPOA = magnocellular preoptic area, SC = the suprachiasmatic nucleus, Str = striatum, TP = posterior tuberculum, VH = ventral hypothalamus, VP = ventral pallium.

**Table 1.** Summary of main statistical effects for neural induction differences.

|  |  | **df** | **F value** | **p value** |
| --- | --- | --- | --- | --- |
| ***R. imitator*** | group | 1,1731 | 13.83 | 0.0002 |
|  | sex | 1,1731 | 0.64 | 0.4242 |
|  | region | 12,1731 | 87.48 | <0.0001 |
|  | sex*group | 1,1731 | 1.25 | 0.2630 |
|  | group*region | 12,1731 | 6.85 | <0.0001 |
|  | sex*region | 12,1731 | 2.69 | 0.0013 |
|  | sex*group*region | 12,1731 | 1.45 | 0.1344 |
| ***D. tinctorius*** | group | 1,2519 | 9.73 | 0.0018 |
|  | sex | 1,2519 | 1.36 | 0.2443 |
|  | region | 12,2519 | 80.15 | <0.0001 |
|  | sex*group | 1,2519 | 1.76 | 0.1844 |
|  | group*region | 12,2519 | 5.00 | <0.0001 |
|  | sex*region | 12,2519 | 3.39 | <0.0001 |
|  | sex*group*region | 12,2519 | 3.89 | <0.0001 |
| ***O. sylvatica*** | group | 1,557 | 1.02 | 0.3126 |
|  | region | 12,557 | 9.40 | <0.0001 |
|  | group*region | 12,557 | 5.53 | <0.0001 |

Comparing neural activity patterns associated with parental care across species allowed us to identify brain regions important in parental care independent of sex and species (i.e. core parental care brain regions). We observed parallel increases in neural activity in tadpole transporting individuals in two core brain regions across all species: the preoptic area (POA) and the medial pallium (Mp; homolog of the mammalian hippocampus). In the POA, patterns differed by subdivision, with female-specific effects in the magnocellular POA and male-specific effects in the anterior POA (Fig. 1C). We also observed increased neural activity in the Mp of non-caregiving female reproductive partners in both *D. tinctorius* and *R. imitator* (Fig. 1C).

### Gene expression in behaviourally relevant neurons

Following identification of core brain regions active during tadpole transport across species and sexes, we sought to identify behaviourally relevant neuronal types within these regions. We found 25 transcripts with significant log-fold expression enrichment/depletion in the POA and 32 transcripts with significant log-fold expression enrichment/depletion in the Mp, seven of which were overlapping between brain regions (Fig. 2; Table 2). Of the overlapping transcripts, four had log-fold expression differences in the same direction (galanin, prolactin receptor, neuropeptide Y receptor 2, brain specific angiogenesis inhibitor associated protein 2) and three had log-fold expression differences in opposite directions (aquaporin 4, dopamine receptor 1B, leptin receptor) between brain regions (Fig. 2).

**Figure 2.**
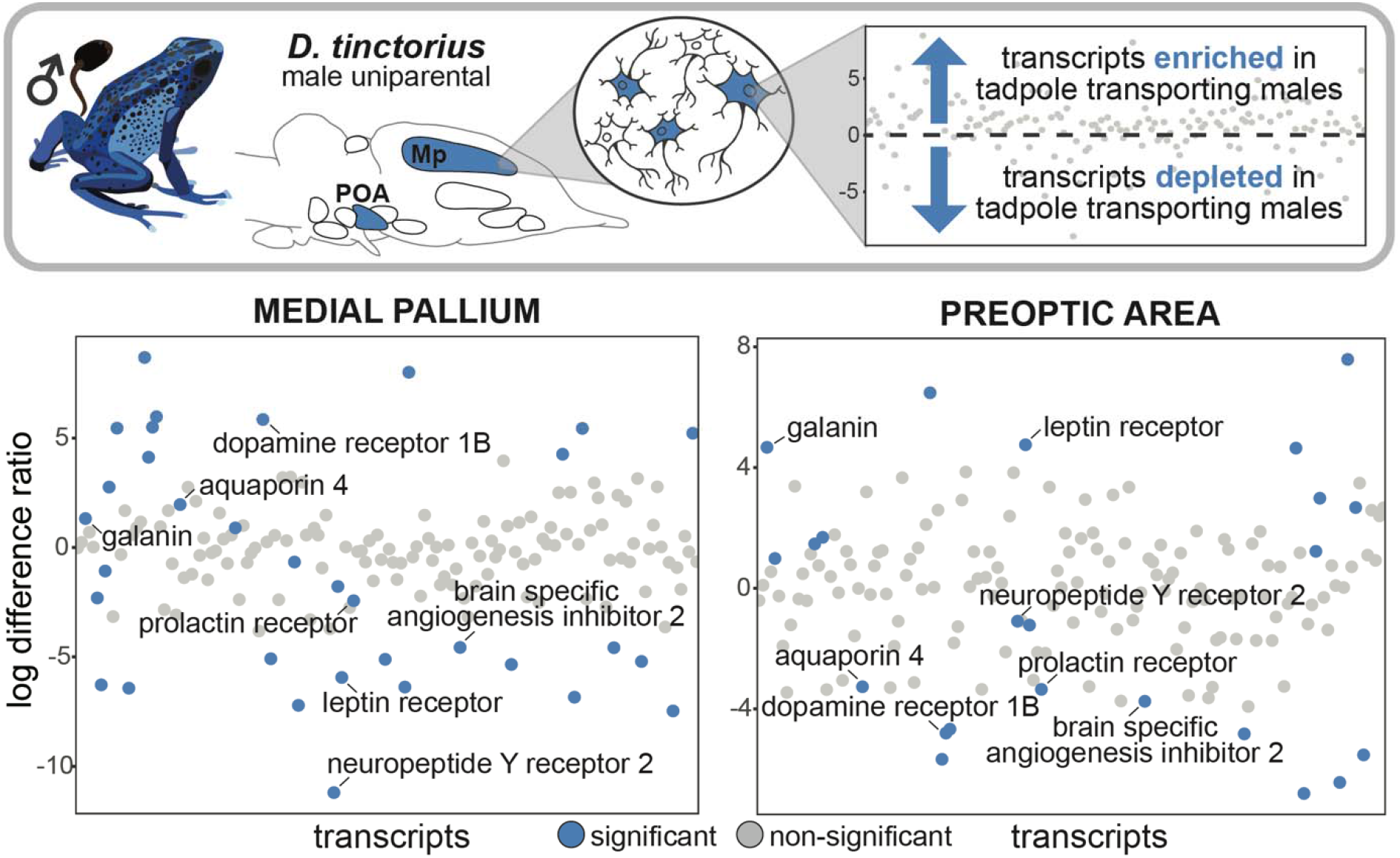
Gene expression in behaviourally relevant neurons. We identified significant expression differences in neurons active during parental care in the preoptic area (POA) and medial pallium (Mp) of tadpole transporting versus non-parental *D. tinctorius* males. We found some unique and some shared transcripts differentially expressed across brain regions (i.e. distribution of blue dots between plots). Those transcripts with significant expression enrichment or expression depletion in tadpole transporting males as compared to control males are highlighted in blue, and the seven transcripts overlapping between brain regions are labeled. The same candidate transcripts are plotted in the same order along the x-axis for both brain regions.

**Table 2.**
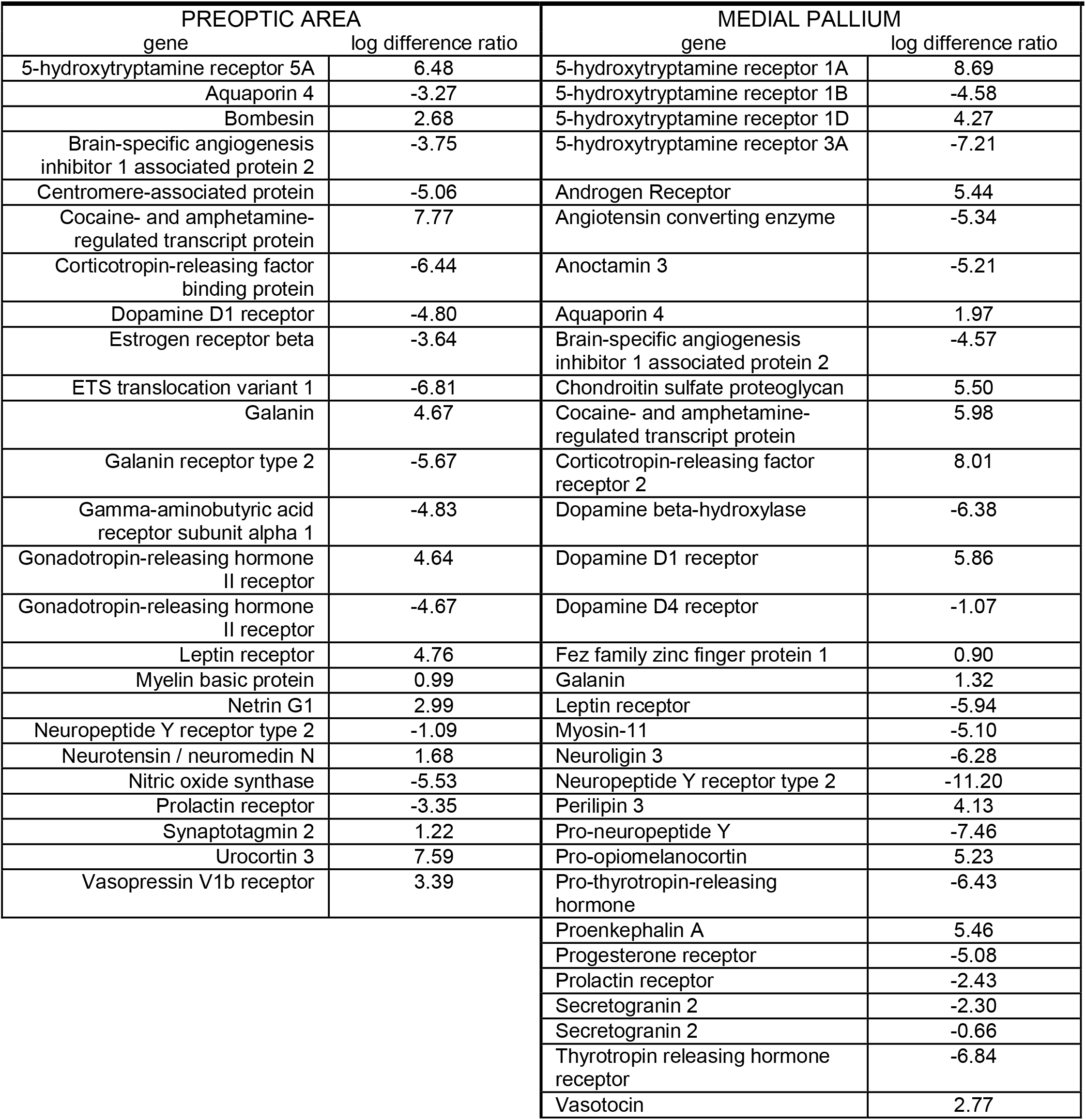
Gene expression in behaviourally relevant neurons. Summary of transcripts significantly enriched (log different ratio >0) or depleted (log difference ratio <0) in tadpole transporting as compared to non-parental male D. tinctorius in the preoptic area (POA) and medial pallium (Mp).

### Galanin neuron number and activity

Recent demonstrations of the importance of galanin in mediating parental care in mice [20,21] and our own findings of galanin transcript enrichment in neurons active during tadpole transport led us to ask whether activity differences specifically in POA galanin neurons were associated with parental care. Parental *R. imitator* had significantly more galanin neurons than did non-parental *R. imitator*, independent of sex (behavioural group: F_1,404_=4.58, p=0.0329), but there were no differences in galanin neuron number in *D. tinctorius* or *O. sylvatica* (Fig. S2). Both *D. tinctorius* and *R. imitator* showed differences in galanin neuron activity associated with parental care, but not in the same manner: in *D. tinctorius* the proportion of active galanin neurons was greater in the female partners of non-transporting males than any other group (sex*behavioural group: F_1,40_=12.73, p=0.0010; Fig. 3). In contrast, in *R. imitator* the proportion of active galanin neurons was greater during tadpole transport in both males and females (behavioural group: F_1,26_=8.15, p=0.0083; Fig. 3). We observed no differences in the proportion of active galanin neurons between tadpole-transporting and non-transporting *O. sylvatica* females.

**Figure 3.**
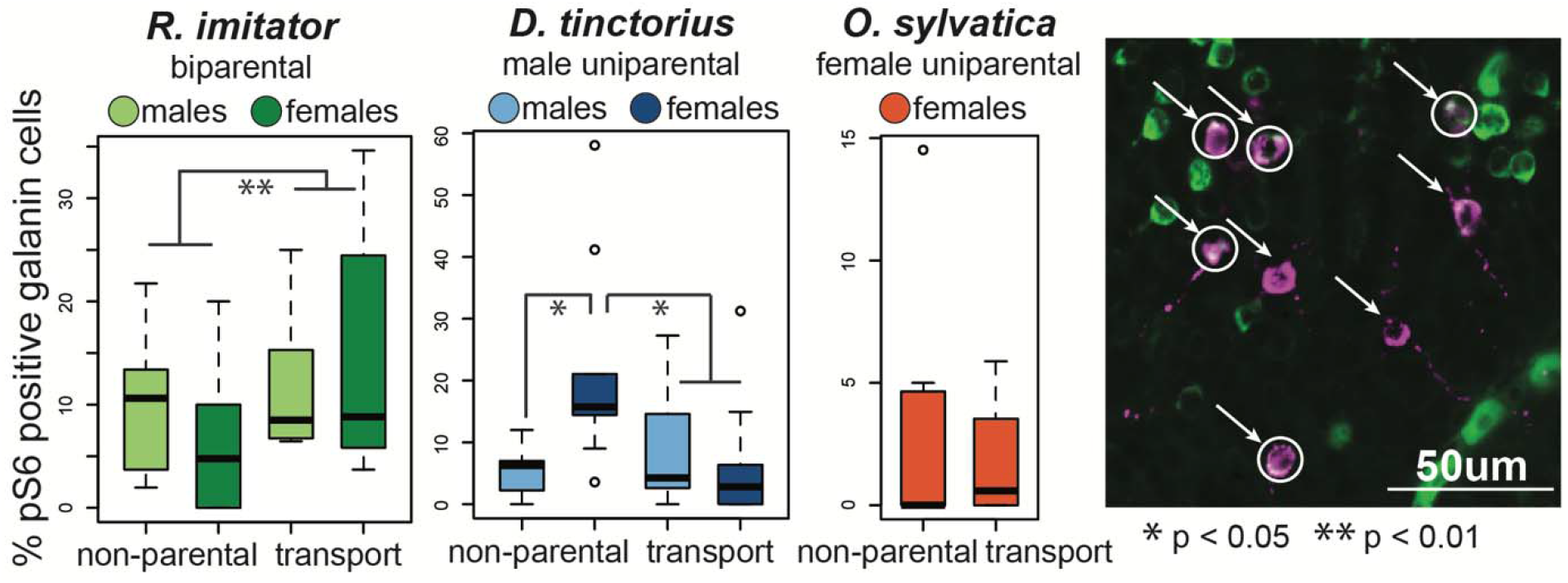
Preoptic area galanin neuron activity. Parental *R. imitator* had a greater proportion of active galanin neurons, as did the female partners of non-parental *D. tinctorius*. Representative micrograph: magenta = galanin positive neurons (arrows), green = pS6 positive neurons, white = co-localization indicating active galanin neurons (circles).

## Discussion

Parental care requires the coordination of hormonal, neural, and molecular changes, many of which remain poorly understood. We took advantage of shared parental behaviour across three poison frog species with distinct parental care strategies, combining lab and field data to disentangle sex- and species-specific mechanisms from core neural mechanisms. We identified the medial pallium and preoptic area as core brain regions associated with parental care and demonstrated expression changes in genes associated with parental care in other vertebrates. Mechanistic studies in closely related, behaviourally variable poison frogs offer a unique opportunity to distinguish shared principles and neural diversity in the mechanisms mediating parental care.

### Core brain regions for parental care

By comparing patterns of neural activity across closely related species with distinct care strategies, we were able to identify core brain regions in which increased neural induction during parental care was sex and species independent. We observed increased neural induction in the medial pallium (Mp) and one or more subdivisions of the preoptic area (POA) during parental care in all focal species. The POA’s widespread connections with other brain regions and high density of neuromodulators make it ideally positioned to modulate complex social behaviour, including parental care. Although data outside mammals is sparse, POA activity is associated with parental behaviour across vertebrates, including mammals [3], birds [3,22], fish [23], and now frogs. In brief, the POA appears to be a core node in parental care circuitry across vertebrates. Importantly, parental care has evolved independently across these clades, indicating convergence across behavioural and neural levels.

Although the precise function of the hippocampus and its non-mammalian homologs remains an area of active research, this brain region is classically implicated in memory, and specifically spatial memory [24,25]. Poison frogs inhabit complex rain forest environments in which tadpole deposition sites are a limited resource of variable quality. Behavioural studies in poison frogs document the use of cognitive spatial maps [26] and demonstrate the importance of spatial memory for navigating back to high-quality tadpole deposition pools [27] and for relocating offspring in egg provisioning species [28]. Increased neural induction in the Mp during tadpole transport is therefore in line with the unique ecological and evolutionary pressures associated with parental care in poison frogs. Indeed, spatial cognition is an important, but rarely examined, component of parental care [29,30], and motherhood is associated with changes in hippocampal plasticity in rodents (reviewed in [31]). Comparisons of hippocampal involvement in parental care across species may yield particularly interesting results given the functional – but not anatomical – conservation of this structure across vertebrates [32].

### Shared parental care circuitry across sexes

The strength of our design is highlighted by identification of inter-specific variation in neural activity patterns between sexes that provide exciting, mechanistic hypotheses to be tested by future studies. Patterns of neural activity during tadpole transport differed between males and females in uniparental *D. tinctorius*, but not biparental, pair bonding *R. imitator*. Females are not directly involved in tadpole transport in either species; however, biparental *R. imitator* females provide parental care in the form of egg attendance prior to tadpole transport and tadpole provisioning following transport [13,32,33]. Thus, similar patterns of neural activity in male and female *R. imitator* could arise either because both sexes are in a “parental state” that modulates long-term circuit activity or because even indirect involvement in tadpole transport activates parental circuitry (i.e. female frogs have to know where their tadpoles are transported in order to return to feed them). In either case, similarities in neural activity patterns associated with parental care in males and females suggest that parental care circuitry is conserved across sexes.

In addition to broad similarities in *R. imitator*, we also observed increased neural activity in the Mp of non-caregiving *D. tinctorius* females. While they are not the typically caregiving sex and do not appear to pair bond with a single male partner, females of *D. tinctorius* and related species will occasionally perform tadpole transport [34,35]. This behavioural flexibility demonstrates that parental circuits are present and can be activated under certain circumstances in females. We suggest an increase in Mp neural activity is related to females’ monitoring of their partners’ behaviour (even in the absence of increased or preferential behavioural affiliation) and ability to perform tadpole transport in the absence of their male partners. In other words, females may monitor male behaviour in order to assess when and if they need to take over parental duties to ensure the survival of their offspring. The diversity of behavioural care strategies between species combined with behavioural flexibility within species in poison frogs affords a unique opportunity to further disentangle the evolution of sex-specific parental care circuits in future.

### Expression variation in behaviourally relevant neurons

Using *D. tinctorius* males, we characterized gene expression differences specifically in neurons active within the POA and Mp during parental care, focusing our analyses on genes previously identified as markers of neuronal types involved in parental care [36]. Of particular interest in the POA were increased expression of the vasopressin 1b receptor, a gonadotropin-releasing hormone receptor, and a number of stress response related genes (Urocortin-3, CART, CRF binding protein) (Table 2). Links between vasopressin and parental care have been demonstrated in rodents [37,38] and vasopressin and gonadotropin releasing hormone may also influence parental care indirectly through their regulation of other molecules with known roles in parental care (e.g. oxytocin, prolactin) [3]. Stress hormones are known to increase in response to the behavioural and metabolic demands of parental care [39,40] providing a link between parental behaviour and the observed upregulation of stress-related signaling pathways.

Notable in the Mp were increased expression of vasopressin and androgen receptor transcripts. As described above, vasopressin signaling is widely implicated in parental care, and has been specifically linked to space use and behavioural and life-history trade-offs in parental prairie voles [29,30]. Space use and navigational abilities differ between males and females in many species, and it has been proposed that greater navigational abilities in males are a side effect of increased androgen signaling [41]. Increased androgen signaling during parental care in poison frogs could facilitate the heightened spatial cognition important during tadpole transport. Furthermore, increasing signaling via region specific receptor expression overcomes the lower testosterone levels typically observed in parental males [42].

In addition to changes specific to either the POA or Mp, we observed a number of transcripts with significant expression differences in both regions. Among them were dopamine and prolactin receptors, and a number of molecules and receptors most commonly implicated in feeding behaviour (galanin, leptin receptor, NPY receptor). Dopamine and prolactin play known roles in parental care [42–44], while other shared transcripts (and some of those unique to a single brain region) are traditionally associated with feeding behaviour. There is growing recognition that molecules traditionally classified as feeding-related play important roles in mediating social behaviour, providing exciting opportunities to explore the repeated targeting of feeding related mechanisms in the convergent evolution of parental care [45].

### Galanin and parental care

Initially described in relation to feeding behaviour, recent work uncovered a role for POA galanin neurons in driving parental care in both male and female mice [20,21]. We found a positive association between parental care and galanin neuron number and activity in biparental *R. imitator*, but not in male uniparental *D. tinctorius*, nor female uniparental *O. sylvatica*. Indeed, the only significant difference outside *R. imitator*, was a relative increase in galanin neuron activity in the female partners of non-transporting male *D. tinctorius*, and we note that the percent of active galanin neurons was overall low in all species.

While recent work demonstrates a sex-independent, behaviour-specific link between galanin neuron activity and parental care [20,21], the earliest work on POA galanin in rodents showed that microinjection of galanin into the POA of male rats facilitated copulatory behaviour [46], and work in fish similarly suggests an association between male courtship behaviour and galanin signaling [47,48]. Thus, species in which the role of galanin in social behaviour has been explored vary in parental care strategy: rats are female uniparental, only some male mice exhibit male care, and fish include both male uniparental and female uniparental species. Together with our findings across frog species with distinct care patterns, these observations suggest that the role of galanin signaling in parental care may be mediated – both acutely and evolutionarily – by life history differences related to parental care, interactions among partners, and male courtship strategy. In brief, galanin appears to have been repeatedly evolutionarily coopted to modulate social behaviour, but the type(s) of social behaviour influenced by galanin signaling are complex, mediated by behavioural variation and evolutionary history, and providing fertile ground for future comparative research.

### Conclusions

Our findings lay the foundation for exciting work using poison frogs as a model to explore neural and molecular mechanisms of parental care, sex-specific behavioural patterns, and the integration of social and environmental cues to coordinate complex social behaviour. We identified core brain regions associated with tadpole transport across dendrobatid poison frogs with distinct care strategies. Moreover, we confirmed a role in amphibians for hormones and neuropeptides associated with parental care in other vertebrates. While increased POA activity was associated with parental care across species, activity specifically of galanin neurons differed between species, suggesting that shared brain regions may nonetheless rely on unique neuronal types to mediate similar behaviour. Studies in closely related but behaviourally distinct species across the animal kingdom provide opportunities to build a more holistic understanding of how shared principles and species-specific diversity govern parental behaviour.

## Supporting information

representative SAS statistical code

phosphoTRAP-Rcode

raw-data-tables

D. tinctorius brain atlas

## Acknowledgements

We thank the O’Connell Lab frog caretakers for help with animal care, Lola Guarderas (Wikiri) and Manuel Morales-Mite (Centro Jambatu) for field work support, and Julie Butler, Hans Hofmann, and the members of the O’Connell Lab for comments on previous versions of the manuscript. We thank Allen Moore and one anonymous reviewer for feedback that improved the manuscript during the review process.

## Data availability

Cell counts, read counts from phosphoTRAP, R code for phosphoTRAP analysis, and the *D. tinctorius* brain atlas are available as Supplemental Materials associated with the manuscript. Raw sequencing reads are available through the NCBI SRA repository (SUB5832932).

## Authors’ contributions

LAO conceived of the study with input from KS and LAC; LAO, EKF, ABR, NAM, and EET collected samples; EKF and LAO performed molecular work and data analysis; EKF and LAO wrote the manuscript with input from all authors. All authors gave final approval for publication and agree to be held accountable for the work performed therein.

