## Supplemental Tables & Figures for "The neural basis of tadpole transport in poison frogs"

**Supplemental Materials**

**Methods**

*Immunohistochemistry*

Whole brains were placed into 4% paraformaldehyde at 4C overnight and then transferred to a 30% sucrose solution for dehydration. Once dehydrated, brains were embedded in Tissue-Tek® O.C.T. Compound (Electron Microscopy Sciences, Hatfield, PA, USA), rapidly frozen, and stored at -80 °C until cryosectioning. We sectioned brains into four coronal series at 14μm, allowed slides to dry completely, and stored slides at -80 °C.

To assess the level of neural activity across brain regions, we used an antibody for phosphorylated ribosomes (pS6; phosphor-S6 Ser235/236; Cell Signaling, Danvers, MA, USA). We followed standard immunohistochemical procedures for 3',3'-diaminobenzadine (DAB) antibody staining. Briefly, we quenched endogenous peroxidases using a 30% sodium hydroxide solution, blocked slides in 5% normal goat serum to reduce background staining, incubated slides in primary antibody (rabbit anti-pS6 at 1:500 in blocking solution) overnight, incubated slides in secondary antibody for 2 hours, followed by avidin-biotin complex (Vectastain Elite ABC Kit; Vector Laboratories, Burlingame, CA, USA) solution incubation for two hours, and treatment with DAB (Vector Laboratories) for 2 min. We rinsed slides with 1X PBS before and after all of the above steps. Finally, slides were rinsed in water, counterstained with cresyl violet, dehydrated in a series of ethanol baths (50%, 75%, 95%, 100%, 100%), and cleared with xylenes prior to cover slipping with permount.

To ask whether neural activity was higher specifically in galanin neurons, we combined the pS6 antibody with a custom-made galanin antibody (peptide sequence: CGWTLNSAGYLLGPHAVDNHRSFNDKHGLA; Pocono Rabbit Farm & Laboratory, Inc, Canadensis, PA, USA). We followed standard immunohistochemical procedures for fluorescent double antibody labeling. Briefly, we blocked slides in 5% normal goat serum to reduce background staining, incubated slides in both primary antibodies (rabbit anti-pS6 at 1:500 and guinea pig anti-galanin at 1:1000 in blocking solution) overnight, and incubated slides in a mix of fluorescent secondary antibodies (AlexaFlour 488 anti-rabbit and AlexaFlour 594 anti-guinea pig at 1:200 in blocking solution) for 2 hours. We rinsed slides with 1X PBS before and after all of the above steps and rinsed slides in water prior to cover slipping using Vectashield with DAPI.

*Phosphotrap library construction*

PhosphoTRAP was performed following Knight et al. (2012) with a few modifications described here. Frozen tissue punches were combined into homogenization buffer (Knight et al., 2012) in 2mL BeadBug™ homogenization tubes (Sigma-Aldrich, St Louis, MO, USA) and homogenized in a BeadBug homogenizer for 3 min at 4 °C. Homogenized lysate was spun for 2000xg at 4 °C at 10m and supernatant was moved to a new tube. After treatment with 10% NP40 and DHPC to clear the supernatant, a 50 uL aliquot was removed for total RNA (TOT) and stored at -80 °C. The remainder of the cleared lysate was incubated with Protein A Dynabeads (Invitrogen, Carlsbad, CA, USA) loaded with the pS6 antibody (phospho-S6 Ser244/247; Invitrogen, Carlsbad, CA, USA) for 10 min at 4 °C and then washed four times with cold 0.35M KCl wash buffer. This immunoprecipitated (IP) RNA was resuspended in buffer RLT from the Qiagen RNeasy Micro kit. Both the IP and total RNA were purified with the RNeasy Micro kit before amplification with the SMART-Seq v4 Ultra Low Input RNA Kit for Sequencing (Takara, Mountain View, CA, USA) and library preparation with the Nextera XT DNA Library Prep kit (Illumina, San Diego, CA, USA), both according to manufactures' instructions. Libraries were then pooled in equimolar amounts and sequenced on an Illumina HiSeq 2500.

*Phosphotrap analysis*

To analyze phosphotrap data we first quantified gene expression in each sample by mapping sequenced reads back to a brain tissue specific *D. tinctorius* de novo transcriptome we previously constructed (Fischer & O'Connell, *unpublished*). Read mapping and quantification were performed using Kallisto software (Bray et al., 2016). As gene expression is known to differ across brain regions (e.g. Lein et al., 2007), and because we were interested primarily in expression differences within brain regions, we performed all subsequent analysis steps separately for the Mp and POA. We removed all transcripts that had <1 transcripts per million in all samples and normalized read counts using DESeq2 in R (Love et al., 2014). We quantified relative gene expression differences in active neurons as a log-fold difference by dividing transcript counts from immunoprecipitated (IP) mRNA by transcript counts from total (TOT) mRNA for each sample and taking the log of these values. Due to the impossibility of zero division, we replaced any zero counts with one prior to this step. Finally, we calculated differential fold enrichment between parental versus non-parental groups by dividing the mean log-fold difference of tadpole transporting individuals by the mean log-fold difference in non-parental animals.

Our primary objective was to utilize phosphoTRAP data to identify Mp and POA cell types whose activity differed between parental and non-parental individuals. To this end we restricted further analysis to a subset of 158 transcripts of interest (Table S3). Transcripts of interest included those annotated as genes with known roles in parental care and/or identified in the only previous study linking cell type to parental care of which we are aware (Moffitt et al., 2018). Although we analyzed brain regions separately, we used the same set of transcripts of interest for both regions.

We inspected differential fold enrichment scores to identify transcripts with large log-fold group differences (differential fold enrichment > 4) and additionally used permutation testing to identify significant differences among scores. Briefly, we generated 250 permuted datasets by randomly re-assigning sample identity to log-fold differences. In order to keep the within sample relationships between IP and TOT mRNA constant, we permuted log-fold differences rather than raw values. From these 250 permuted datasets we calculated t-values for the difference between tadpole transport and non-transport groups and generated distributions of these t-values for each transcript. We calculated t-values in an identical manner for the real data and then compared real t-values to the distributions generated from the permuted datasets. For each transcript, if the real t-value fell in the extreme 1% at either end of the permuted t-value distribution we called the transcript differentially expressed between behavioral groups. Permutation testing and visualization were done using R Statistical Software (version 3.5.0; the R Foundation for Statistical Computing).

### Tables

**Table S1.** Genes of interest targeted in phosphoTRAP analysis. Submitted as excel file.

**Table S2.** Posthoc statistical results of tests for group*sex (*D. tinctorius* and *R. imitator*) and group (*O. sylvatica*) differences by brain region.

| | region | df | F value | p value |
|---|---|---|---|---|
| ***R. imitator*** | Acc | 3,1731 | 2.61 | 0.0502 |
| | BST | 3,1731 | 5.90 | **0.0005** |
| | DV/VP | 3,1731 | 5.30 | **0.0012** |
| | H | 3,1731 | 1.73 | 0.1598 |
| | Ls | 3,1731 | 10.87 | **<.0001** |
| | Mgv | 3,1731 | 4.68 | **0.0029** |
| | Mp | 3,1731 | 9.75 | **<.0001** |
| | aPOA | 3,1731 | 7.14 | **<.0001** |
| | mPOA | 3,1731 | 5.86 | **0.0006** |
| | SC | 3,1731 | 4.51 | **0.0037** |
| | Str | 3,1731 | 2.15 | 0.0924 |
| | TP | 3,1731 | 0.10 | 0.9599 |
| | VH | 3,1731 | 1.03 | 0.3795 |
| ***D. tinctorius*** | Acc | 3,2519 | 4.82 | **0.0024** |
| | BST | 3,2519 | 0.51 | 0.6740 |
| | DV/VP | 3,2519 | 1.28 | 0.2808 |
| | H | 3,2519 | 4.61 | **0.0032** |
| | Ls | 3,2519 | 11.72 | **<.0001** |
| | Mgv | 3,2519 | 2.03 | 0.1082 |
| | Mp | 3,2519 | 9.85 | **<.0001** |
| | aPOA | 3,2519 | 4.16 | **0.0060** |
| | mPOA | 3,2519 | 1.83 | 0.1390 |
| | SC | 3,2519 | 2.86 | **0.0355** |
| | Str | 3,2519 | 2.53 | 0.0557 |
| | TP | 3,2519 | 4.91 | **0.0021** |
| | VH | 3,2519 | 2.71 | **0.0434** |
| ***O. sylvatica*** | Acc | 1,557 | 0.06 | 0.8065 |
| | BST | 1,557 | 0.52 | 0.4699 |
| | DV/VP | 1,557 | 0.03 | 0.8675 |
| | H | 1,557 | 3.45 | 0.0637 |
| | Ls | 1,557 | 1.38 | 0.2412 |
| | Mgv | 1,557 | 0.24 | 0.6242 |
| | Mp | 1,557 | 13.57 | **0.0003** |
| | aPOA | 1,557 | 046 | 0.4956 |
| | mPOA | 1,557 | 2.99 | 0.0842 |
| | SC | 1,557 | 0.00 | 0.9908 |
| | Str | 1,557 | 2.20 | 0.1382 |
| | TP | 1,557 | 0.03 | 0.8525 |
| | VH | 1,557 | 0.16 | 0.6917 |

**Figures**

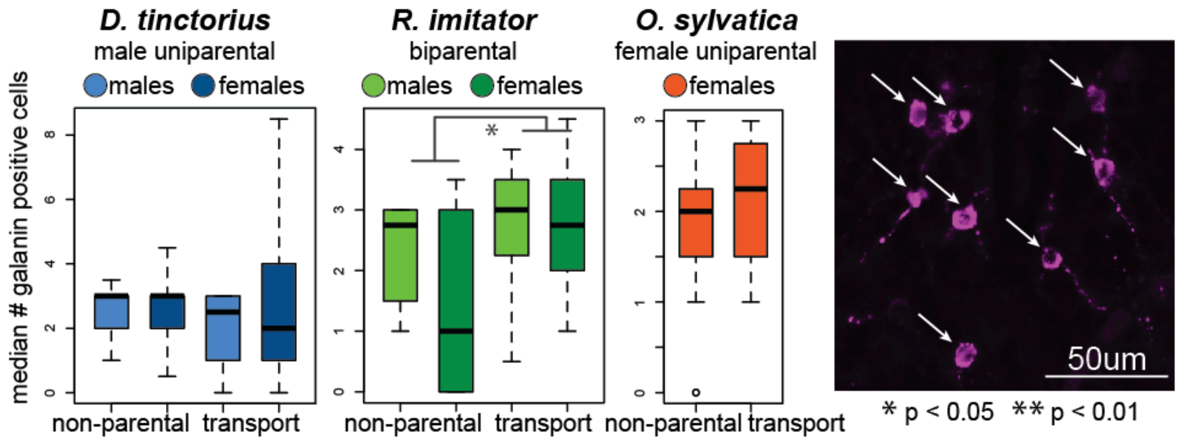


**Figure S1.** Preoptic area galanin neuron number. Parental *R. imitator* had more POA galanin neurons than non-parental R. imitator, regarless of sex. There were no differences in galanin neuron number based on sex or behavioral group in *D. tinctorius* or *O. sylvatica*. Representative micrograph: magenta = galanin positive neurons (arrows).
