## Supplementary material for "The neural basis of tadpole transport in poison frogs": D. tinctorius brain atlas

### NEUROANATOMICAL ATLAS OF THE DYEING POISON FROG (*Dendrobates tinctorius*)

Lauren A O'Connell

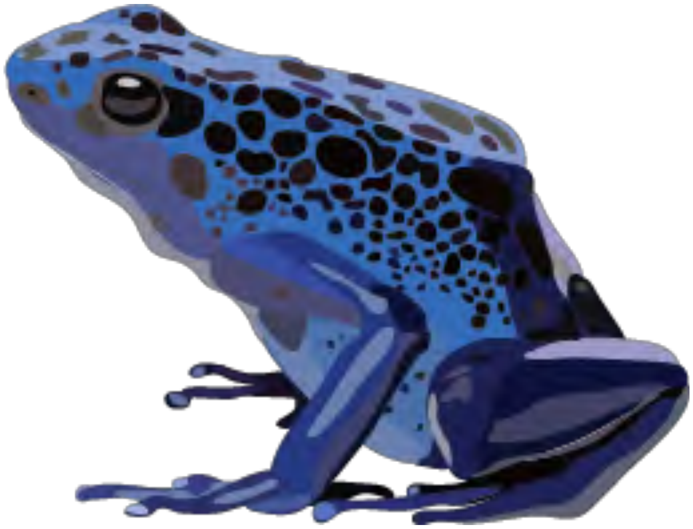

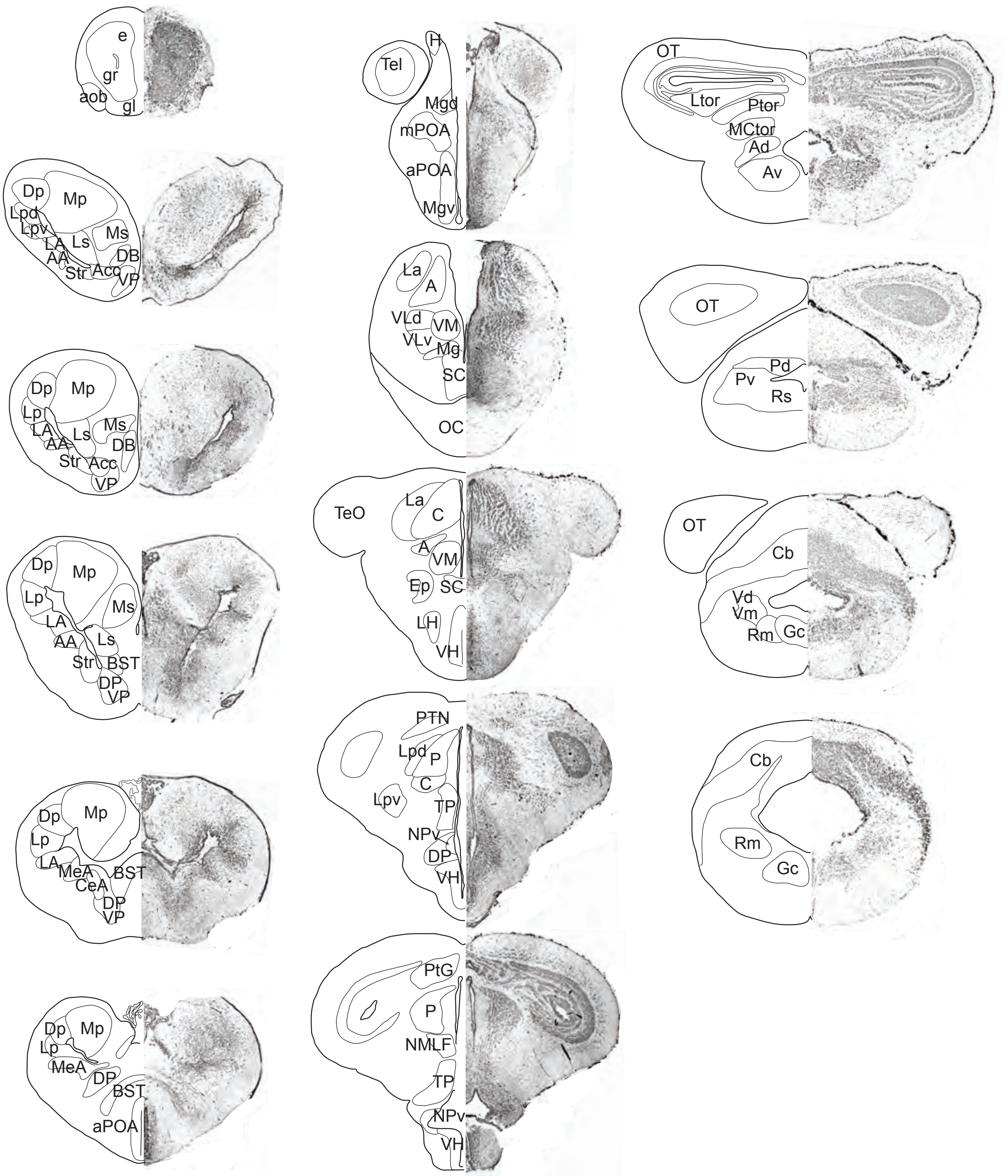

e
gr
aob
gl
Dp
Mp
Lpd
Lpv
Ms
LA
Ls
DB
AA
Str
Acc
VP
Dp
Mp
Lp
Ms
LA
Ls
DB
AA
Str
Acc
VP
Dp
Mp
Lp
Ms
LA
Ls
AA
Str
BST
DP
VP
Dp
Mp
Lp
LA
MeA
BST
CeA
DP
VP
Dp
Mp
Lp
MeA
DP
BST
aPOA
H
Tel
Mgd
mPOA
aPOA
Mgv
La
A
VLd
VM
VLv
Mg
SC
OC
TeO
La
C
A
VM
Ep
SC
LH
VH
PTN
Lpd
P
C
Lpv
TP
NPv
DP
VH
PtG
P
NMLF
TP
NPv
VH
OT
Ltor
Ptor
MCtor
Ad
Av
OT
Pd
Pv
Rs
OT
Cb
Vd
Vm
Rm
Gc
Cb
Rm
Gc

#### List of Abbreviations

| | |
|---|---|
| A | anterior thalamic nucleus |
| AA | anterior amygdaloid area |
| Acc | nucleus accumbens |
| Ad | anterodorsal tegmental nucleus |
| AH | anterior hypothalamus |
| aob | accessory olfactory bulb |
| Av | anteroventral tegmental nucleus |
| BST | bed nucleus of the stria terminalis |
| C | central thalamic nucleus |
| Cb | Cerebellum |
| CeA | central amygdala |
| DB | diagonal band of Broca |
| DH | dorsal hypothalamic nucleus |
| Dp | dorsal pallium |
| DP | dorsal pallidum |
| e | postolfactory eminence |
| Ep | posterior entopeduncular nucleus |
| Gc | griseum centrale rhombencephali |
| gl | glomerular layer of the olfactory bulb |
| gr | granule cell layer of the olfactory bulb |
| Hv | ventral habenula |
| La | lateral thalamic nucleus, anterior division |
| LA | lateral amygdale |
| LH | lateral hypothalamic nucleus |
| Lp | lateral pallium |
| Lpd | lateral thalamic nucleus, posterodorsale |
| Lpv | lateral thalamic nucleus, posteroventrale |
| Ls | lateral septum |
| M | dorsal midline |
| MeA | medial amygdale |
| Mgd | magnocellular preoptic nucleus, dorsal part |
| Mgv | magnocellular preoptic nucleus, ventral part |
| ml | mitral cell layer of the olfactory bulb |
| Mp | medial pallium |
| Ms | medial septum |
| ON | optic nerve |
| Npv | nucleus of the periventricular organ |
| P | posterior thalamic nucleus |
| Pd | nucleus posterodorsalis tegmenti |
| aPOA | anterior preoptic area |
| mPOA | medial preoptic area |
| Pv | nucleus posteroventralis tegmenti |
| Rm | nucleus reticularis medius |
| Rs | nucleus reticularis superior |
| SC | suprachiasmatic nucleus |
| Str | Striatum |
| Tect | optic tectum |
| Tel | telencephalon |
| Tor-L | torus semicircularis, laminar nucleus |
| Tor-P | torus semicircularis, principal nucleus |
| Tor-V | torus semicircularis, ventral area |
| TP | posterior tuberculum |
| Vd | descending trigeminal tract |
| VH | ventral hypothalamic nucleus |
| VLd | ventrolateral thalamic nucleus, dorsal part |
| VLv | ventrolateral thalamic nucleus, ventral part |
| Vm | nucleus motorius nervi trigemini |
| VM | ventromedial thalamic nucleus |
| VP | ventral pallidum |
